# The Src-family kinase Lyn plays a critical role in establishing and maintaining B cell anergy by suppressing PI3K-dependent signaling

**DOI:** 10.1101/2024.05.21.595208

**Authors:** Brigita E. Fiske, Scott M. Wemlinger, Bergren W. Crute, Andrew Getahun

## Abstract

Although the Src family kinase (SFK) Lyn is known to be involved in induction and maintenance of peripheral B cell tolerance, the molecular basis of its action in this context remains unclear. This question has been approached using conventional as well as B cell-targeted knockouts of Lyn, with varied conclusions likely confused by collateral loss of Lyn functions in B cell and myeloid cell development and activation. Here we utilized a system in which Lyn gene deletion is tamoxifen inducible and B cell restricted. This system allows acute elimination of Lyn in B cells without off-target effects. This genetic tool was employed in conjunction with immunoglobulin transgenic mice in which peripheral B cells are autoreactive. DNA reactive Ars/A1 B cells require continuous inhibitory signaling, mediated by the inositol phosphatase SHIP-1 and the tyrosine phosphatase SHP-1, to maintain an unresponsive (anergic) state. Here we show that Ars/A1 B cells require Lyn to establish and maintain B cell unresponsiveness. Lyn primarily functions by restricting PI3K-dependent signaling pathways. This Lyn-dependent mechanism complements the impact of reduced mIgM BCR expression to restrict BCR signaling in Ars/A1 B cells. Our findings suggest that a subset of autoreactive B cells requires Lyn to become anergic and that the autoimmunity associated with dysregulated Lyn function may, in part, be due to an inability of these autoreactive B cells to become tolerized.

## Introduction

The Src family kinase Lyn plays dual roles in B cell activation (1, 2). It is important early in B cell receptor (BCR) signaling, phosphorylating the Immunoreceptor Tyrosine-based Activation Motifs (ITAMs) of CD79A and B, as well as Syk and CD19 (3, 4). Lyn is also important in regulatory signaling, phosphorylating Immunoreceptor Tyrosine-based Inhibition Motifs (ITIMs) on inhibitory receptors, such as FcγRIIB (5–8), when these are brought into proximity of actively signaling BCRs, as well as by additional less well-defined mechanisms (9). In the absence of Lyn, other Src family kinases, such as Fyn, compensate for Lyn’s role in activating BCR signals, but not inhibitory signaling (5, 7, 10). Consequently, B cell-restricted Lyn deficiency results in B cell hyperresponsiveness and a loss of B cell tolerance (11). The importance of Lyn-mediated inhibitory signaling in B cell tolerance is further demonstrated by the observation that besides Lyn deficiency (8, 11–13), deficiencies of upstream inhibitory receptors (e.g. CD22 (14, 15), FcγRIIB (16, 17) and CD72 (18)) as well as deficiencies of downstream effector phosphatases (SHP-1 (19) and SHIP-1 (20, 21)) result in a loss of B cell tolerance and lupus-like autoimmunity. This pathway is often dysregulated in individuals that develop B cell-dependent autoimmunity (22–29).

How Lyn contributes to peripheral B cell tolerance is not fully understood (11, 30, 31). B cell anergy is the most prevalent peripheral B cell tolerance mechanism (32). Anergic B cells exist in a state of functional unresponsiveness in the periphery (33). This anergic state is thought to result from continuous BCR stimulation with self-antigen in the absence of T cell help (34, 35). One of the characteristics of anergic B cells is that they are hyporesponsive to BCR stimulation (36). Both reduced IgM BCR surface expression (37, 38) and inhibitory signaling mechanisms (39, 40) contribute to this state of hyporesponsiveness, although the precise contribution of each mechanism is unclear. Since Lyn plays a critical role in establishing inhibitory signaling circuits in B cells, one would expect that Lyn is involved in establishing B cell anergy, but previous studies were inconclusive (30, 31).

To address this question, we used the Ars/A1 model of B cell anergy in conjunction with genetic models that allow us to delete Lyn in a B cell specific and inducible manner. Ars/A1 B cells express a transgenic BCR that binds the hapten p-azophenylarsonate (Ars) but cross-reacts with lower affinity with DNA-containing self-antigens in vivo, rendering Ars/A1 B cells anergic (41). Previously, we demonstrated that Ars/A1 B cells require continuous activity of the inositol phosphatase SHIP-1 and the tyrosine phosphatase SHP-1 to establish and maintain an anergic state (40). Given its dependence on inhibitory signaling, we asked whether this model of B cell anergy, representing anergic B cells that depend strongly on inhibitory signaling, requires Lyn to become anergic. Indeed, we found that Ars/A1 B cells require Lyn to establish and maintain an anergic state. Mechanistically, this is due to Lyn-dependent restriction of PI3K-dependent signaling. Impaired activation of PI3K-independent signaling events in Ars/A1 B cells is primarily a consequence of strongly reduced IgM antigen receptor expression. Taken together these findings show that Lyn plays an important role in establishing an anergic state in a subset of autoreactive B cells. It further suggests that the development of autoimmune disease associated with dysregulation of Lyn function may in part be due to compromised anergic B cells.

## Material and Methods

### Mice

Six- to twenty-week-old mice were used in all experiments. In each experiment, mice were age-matched and sex-matched. Both sexes were used in our experiments with similar results. As described previously (40), hCD20-TamCre animals (42) were intercrossed with mice carrying the rosa26-flox-STOP-YFP allele (43), generating mice in which YFP is expressed in B cells upon Cre activation. These mice were crossed with Ars/A1 (41) and MD4 (44) B cell antigen receptor transgenic mice to generate hCD20-TamCre x rosa26-flox-STOP-YFP x Ars/A1 mice and hCD20-TamCre x rosa26-flox-STOP-YFP x MD4 mice, respectively. B cells from these mice will be referred to as WT Ars/A1 and WT MD4. To generate mice in which Lyn deletion can be induced in B cells, these mice were crossed with Lyn^flox/flox^ mice (donated by Dr. C. Lowell, UCSF, San Francisco, CA) (45). To generate mice in which SHIP-1 and Lyn haploinsufficiency can be induced in Ars/A1 B cells, these mice were crossed with previously described mice (40) in which SHIP-1 deletion can be induced in B cells (hCD20-TamCre x rosa26-flox-STOP-YFP x SHIP-1^flox/flox^ x Ars/A1 mice). In parallel, lines were generated that do not express a transgenic BCR to determine the effect of acute Lyn deletion in B cells from mice that express a WT repertoire of BCRs. Mice in which Lyn deficiency is constitutively restricted to the B cell lineage from early B cell development were generated by crossing Lyn^flox/flox^ mice with mb1cre mice (46). This line was crossed with Ars/A1 mice to generate mb1cre x Lyn^flox/flox^ x Ars/A1 mice. Lyn^flox/flox^ x Ars/A1 mice littermates were used as Lyn sufficient controls.

In adoptive transfer experiments, C57BL/6J mice were used as recipients. For some of the signaling experiments, Ars/A1 and MD4 B cell antigen receptor transgenic mice were used without additional strains crossed in. Mice were housed and bred at the University of Colorado Denver (UCD) Anschutz Medical Campus Vivarium (Aurora, CO), with the exception of C57BL/6J mice which were purchased from Jackson Laboratories. All experiments with mice were performed in accordance with the regulations and with approval of the University of Colorado SOM Institutional Animal Care and Use Committee.

### Tamoxifen treatments, adoptive transfers and immunizations

Cre activity was induced by tamoxifen treatment. Tamoxifen (Sigma, T-5648) was dissolved in 10% ethanol (Decon Laboratories) and 90% corn oil (Sigma) at 20 mg/mL. In experiments in which we analyzed the impact of induced Lyn deletion on BCR signaling of splenic B cells, mice were injected once i.p. with 100 μL (2 mg) of tamoxifen/corn oil 7 days prior to analysis.

In adoptive transfer experiments in which we analyzed the impact of induced Lyn deletion on the ability of Ars/A1 B cells to maintain B cell tolerance, recipient mice were irradiated with 200 rads four hours prior to adoptive transfer to provide space for transferred cells as described previously (40). B cells from donor mice were enriched by depletion of CD43^+^ cells with anti-CD43-conjugated magnetic beads (CD43 (Ly-48), mouse; Miltenyi Biotec). Resultant populations were >97% B cells based on B220 staining and FACS analysis. Prior to transfer, donor B cells were labeled with CellTrace Violet Proliferation Kit, *for flow cytometry* (Invitrogen by Thermo Fisher) at 5 μM for 3 min at RT in complete medium (IMDM supplemented with 10% fetal calf serum (FCS), 1 mM Sodium Pyruvate, 2 mM L-Glutamine, 100 U/mL Pen/Strep, 50 mg/mL gentamicin and 0.1 mM 2-Me). The labeling was stopped by adding an excess of FCS, followed by 3 washes with PBS. 1-8×10^6^ B cells in 200 μL PBS were adoptively transferred by i.v. injection. Twenty-four hours after transfer, Cre activity was induced by 100 μL (2 mg) tamoxifen injections i.p. on two consecutive days.

In experiments in which we analyzed the impact of induced Lyn deletion on the ability of MD4 B cells to mount an antibody response to SRBC-HEL immunization, donor hCD20-TamCre x rosa26-flox-STOP-YFP x Lyn^flox/flox^ x MD4 mice and hCD20-TamCre x rosa26-flox-STOP-YFP x MD4 mice were injected i.p. with 100 μL (2 mg) of tamoxifen/corn oil. Six days post tamoxifen treatment, splenocytes were stained with B220-Alexa647 (clone RA3-6B2, Biolegend) and B220+ YFP+ cells were sorted using a Beckman Coulter Astrios EQ cell sorter. 1.3×10^5^ YFP+ B220+ cells were adoptively transferred by i.v. injection in 200 μL PBS into C57BL/6 recipient mice. Twenty-four hours post adoptive transfer the recipient mice were immunized by i.p. injection with 200 μL 1% SRBC-HEL. Sheep red blood cells (SRBC) were purchased from the Colorado Serum Company and stored in sterile Alsever’s solution at 4 °C. Hen Egg Lysozyme (HEL) was chemically coupled to SRBC. One mL packed SRBC was resuspended in 15 mL PBS containing 5 mg/mL HEL (Sigma). To cross-link, 1mL of 50 mg/mL 1-ethyl-3-(3-dimethylaminopropyl)carbodiimide hydrochloride (Sigma) in PBS was added, mixed and incubated for 1 h at RT, with occasional mixing. Afterwards, the cells were washed four times in PBS and the conjugation of HEL to the SRBC cell surface was confirmed by flow cytometry after staining with Dylight650 conjugated anti-HEL (clone D1.3, produced and conjugated in house). Five days post immunization splenocytes were harvested and analyzed for MD4 anti-HEL responses.

### ELISPOT

For detection of IgM^a^ Ars antibody secreting cells, microtiter plates were coated with 10 μg/mL Ars-BSA_16_ or 10 μg/mL Ars-OVA_8_ in PBS. For detection of IgM^a^ MD4 antibody secreting cells, microtiter plates were coated with 10 μg/mL HEL in PBS. Before use, the plates were washed twice with PBS-0.05% Tween-20 and blocked with complete 10% FCS IMDM medium. Two-fold serial dilutions were made of splenic single cell suspensions starting at 1/50th of a spleen in the first well. The plates were incubated at 37°C at 7% CO_2_ overnight. MD4 and Ars/A1-derived antibody secreting cells were detected with Biotin-conjugated Mouse Anti-Mouse IgM[a] (Clone:DS-1; BD Biosciences), followed by Streptavidin-AP (SouthernBiotech). Between each step, the plates were washed 4 times with PBS-0.05% Tween-20. The plates were developed by incubating with ELISPOT development buffer (25 μM 5-bromo-chloro-3-indolyl phosphate p-toluidine, 100 mM NaCl, 100 mM Tris, 10 mM MgCl_2_ [pH 9.5]) for 1 h. The reaction was stopped by washing the plate once with double-distilled H_2_O. The number of spots at a cell dilution in the linear range was determined, and the number of antibody-secreting cells per spleen was calculated.

### Flow cytometry

Single cell suspensions of splenic cells or bone marrow cells, flushed out of the two hindleg femurs, were prepared in 10% FCS complete medium, and red blood cells were lysed using Ammonium Chloride Potassium (ACK) Lysis buffer (15 mM NH_4_Cl, 10 mM KHCO_3_, 0.1 mM Na_2_EDTA, pH 7.4). Cells were resuspended in PBS containing 1% BSA and 0.05% sodium azide and incubated with an optimal amount of directly fluorochrome labeled antibodies. Antibodies directed against the following molecules were used: B220 (clone RA3-6B2 Biolegend), CD43 (clone S11, Biolegend), CD69 (clone H1.2F3, BD Biosciences), CD86 (clone GL1, BD Biosciences), CD95 (clone Jo2, BD Biosciences), IgD (clone 11-26c.2a, Biolegend), and IgM (B-7-6). B-7-6 was produced in our own laboratory and was directly conjugated to DyLight (Pierce/Thermo Scientific) fluorochromes, according to the manufacturer’s protocol. The MD4 BCR was stained with its antigen HEL (Sigma) conjugated to Dylight 650 (Pierce/Thermo Scientific) fluorophore, according to the manufacturer’s protocol.

To determine the phenotype of Ars/A1 B cells following adoptive transfer, splenocytes were fixed and permeabilized with BD Cytofix/Cytoperm™ and stained with Dylight650-E4 anti-Ars/A1 idiotype (E4 was produced and conjugated to DyLight™ 650 NHS Ester (Pierce/Thermo Scientific) in our laboratory), Dylight488 anti-GFP (polyclonal Goat anti-GFP Antibody (600-101-215, Rockland) conjugated to DyLight™ 488 NHS Ester (Pierce/Thermo Scientific) in our laboratory), and CD138-PE-Cy7 (Clone: 281-2; Biolegend). The cells were washed twice, and events were collected on a CyAn ADP (Dakocytomation), a BD LSRFortessa™ X-20 Cell Analyzer (BD Biosciences) or a Cytek Northern Lights (Cytek Biosciences) and analyzed using FlowJo software (Becton Dickinson & Company (BD). In all experiments, the same gating strategy was used. Following doublet exclusion, we gated on the lymphocyte population based on FSC/SSC properties but including larger cells to capture plasmablasts, followed by gating on YFP+ E4+ events. To calculate the absolute number of YFP+ Ars/A1 B cells per spleen in different states (undivided, proliferated and plasmablasts (proliferated and CD138+), we multiplied the total number of splenocytes (counted with a Beckman Coulter Vi-CELL) with the frequency of YFP+ E4+ events and the frequency of violet proliferation dye^hi^ CD138^lo^ (undivided), violet proliferation dye^lo^ (proliferated), violet proliferation dye^lo^ CD138^hi^ (plasmablasts).

### Analysis of BCR signaling by flow cytometry

To analyze protein phosphorylation by flow cytometry, splenocytes were resuspended in serum-free IMDM at a concentration of 10^7^ cells/mL and incubated at 37°C for 20-30 minutes prior to stimulation. In some experiments the PI3K inhibitor Ly294002 (Cell Signaling Technology), dissolved in DMSO, was added at 5 μM during the last 10 minutes of prewarming. Control tubes received DMSO only at an equal dilution. In some experiments the splenocytes were prestained with a Fab fragment of B-7-6 conjugated to Dylight 488 at 4 °C and washed twice with serum free IMDM prior to incubation at 37 °C to allow for gating on subpopulations of B cells with equal IgM expression. B cells were stimulated with 10 μg/mL polyclonal F(ab’)_2_ goat anti-mouse IgM (anti-μ) (Jackson ImmunoResearch) or 5 μg/mL anti-IgD^a^ (clone AMS 9.1, Santa Cruz Biotechnology) for the indicated time. Prior to stimulation with anti-IgD^a^ the splenocytes were incubated with 10 μg/mL clone 2.4G2 to block binding to FcγRIIB.

Signaling was stopped by adding formaldehyde (Tousimis) at a final concentration of 2%. If a methanol permeabilization method was used the cells were incubated at 37°C for 10 min with the 2% formaldehyde. Afterwards the cells were spun down and resuspended in −80 °C 100% MeOH (Fisher Scientific) and incubated on ice for 30 min. After permeabilization the cells were washed three times in PBS containing 1% BSA. Cell staining and washes were done in PBS/1% BSA. If BD Cytofix/Cytoperm permeabilization was used, the cells were immediately spun down following formaldehyde addition and resuspended in BD Cytofix/Cytoperm. The cells were incubated at 4 °C for 20 minutes, followed by three washes with BD Perm/Wash buffer. Cell staining and washes were done in BD Perm/Wash buffer.

Methanol permeabilized cells were stained for: B220 (clone RA3-6B2 Biolegend), Dylight488 anti-GFP (if cells express YFP), pCD79A (pY182) (clone D189, Cell Signaling Technology), pSyk (pY352) (Clone 17A/P-ZAP70, BD Biosciences), pErk (pT202/pY204) (clone 20A, BD Biosciences), pPLCγ2 (pY759) (clone K86-689.37, BD Biosciences), pBLNK (pY84) (clone J117-1278, BD Biosciences), Rabbit isotype control (Cell Signaling Technology), mouse IgG1 isotype control (clone MOPC-21, BD Biosciences). BD permeabilized cells were stained for: B220 (RA3-6B2 Biolegend), Dylight488 anti-GFP (if cells express YFP), pAkt (pS473) (clone M89-61, BD Biosciences), IκBα (clone L35A5, Cell Signaling Technology), pS6 (pS235/pS236) (clone cupk34k, eBioscience/Invitrogen), Rabbit isotype control (Cell Signaling Technology), mouse IgG1-Alexa647 isotype control (clone MOPC-21, BD Biosciences) and mouse IgG1-PE-Cy7 isotype control (clone P3.6.2.8.1, eBioscience/Invitrogen). The cells were stained for 1hr at RT and washed 3 times prior to analysis. Events were collected on a BD LSRFortessa™ X-20 Cell Analyzer (BD Biosciences) or a Cytek Northern Lights (Cytek Biosciences) and analyzed using FlowJo software (Becton Dickinson & Company (BD). In all experiments, the same gating strategy was used. Following doublet exclusion, we gated on the lymphocyte population based on FSC/SSC properties, followed by gating on YFP+ B220+ events or B220+ events if the cells did not express YFP.

### Analysis of calcium mobilization

For measurements of intracellular free calcium concentration ([Ca^2+^]_i_), splenocytes (2*10^7^/mL in IMDM medium containing 2% FCS) were simultaneously stained with anti-B220-Alexa647 (clone RA3-6B2 Biolegend) and loaded with 5 μM Indo-1 acetoxymethyl (Indo1-AM) (Molecular Probes) for 30 min at RT. In some experiments 5 μM Ly294002 or B-7-6 Fab conjugated to Dylight 488 was added. After washing once with IMDM with 2% FCS, the cells were resuspended at 10^7^ cells/mL in warm IMDM with 2% FCS in a 500μL volume. Indo-1 was excited with a 355 nm laser, Ca^2+^-bound Indo-1 was detected using a 379/28 bandpass filter and unbound Indo-1 was detected using a 450/50-410LP bandpass filter. Relative free calcium concentration was determined by the ratio bound/unbound Indo-1. After the baseline was established by analysis for 30 seconds, cells were stimulated with 2-10 μg/mL F(ab’)_2_ goat anti-mouse IgM (anti-μ) (Jackson ImmunoResearch) or 5 μg/mL anti-IgD^a^ (clone AMS 9.1, Santa Cruz Biotechnology) for 2 minutes. Prior to stimulation with anti-IgD^a^ the splenocytes were incubated with 10 μg/mL clone 2.4G2 to block binding to FcγRIIB. Relative mean [Ca^2+^]_i_ was measured using a BD LSRFortessa™ X-20 Cell Analyzer (BD Biosciences) and analyzed using FlowJo software (Becton Dickinson & Company (BD). The area under the curve was calculated in FlowJo.

### Immunoblotting

Splenic B cells were enriched by depletion of CD43^+^ cells with anti-CD43-conjugated magnetic beads (MACS anti-mouse CD43; Miltenyi Biotec). Resultant populations were >97% B cells based on B220 staining and FACS analysis. B cells were resuspended at 10^7^/mL in serum-free IMDM and incubated at 37 °C for 20-30 minutes. B cells were stimulated with 10 μg/mL F(ab’)_2_ goat anti-mouse IgM (anti-μ) (Jackson ImmunoResearch). At the indicated times, the B cells were pelleted by centrifugation at 6000 rpm for 30 seconds in a tabletop centrifuge and lysed in reducing SDS sample buffer or cells were lysed in 1% NP-40 lysis buffer (25 mM Tris-HCl (pH 7.4), 150 mM NaCl, 1% NP-40, 1 mM EDTA, 1 mM DTT, with protease and phosphatase inhibitors (100 µg/mL aprotinin, 100 µg/mL α-1-antitrypsin, 100 µg/mL leupeptin, 1 mM PMSF, 10 mM NaF, 2 mM NaVO_3_, and 10 mM tetrasodium pyrophosphate). For analysis, whole cell lysates (WCLs) were mixed with a quarter final volume of 4X reducing Laemmli SDS-PAGE sample buffer. To validate Lyn deletion following tamoxifen treatment, seven-day post tamoxifen treatment splenocytes were stained with B220-Alexa647 (clone RA3-6B2, Biolegend) and B220+ YFP+ cells were sorted using a Beckman Coulter Astrios EQ cell sorter, counted and lysed in reducing Laemmli SDS-PAGE sample buffer.

Samples (0.5-1 × 10^6^ cell equivalents) were boiled for 5 minutes and proteins were separated by SDS-PAGE, and transferred to PVDF membranes, which were then blocked with 5% BSA in TBST (TBS-0.05% tween-20). Blots were probed with antibodies diluted in 3% BSA-TBST. Antibodies specific for pAkt (S473) (Cell Signaling), p44/42 MAPK (Erk1/2) (Thr202/Tyr204) (Cell Signaling Technology), pan Erk (BD), Akt (Cell Signaling), pSyk (Tyr525/Tyr526) (Cell Signaling Technology), Syk (Cell Signaling), Lyn (affinity purified polyclonal rabbit IgG raised against Lyn 1-131) or actin (clone C-2, Santa Cruz) were used. Following incubation with primary antibodies, the blots were washed in TBS-0.05% tween-20 and incubated with a secondary antibody HRP-conjugated Goat anti-Rabbit IgG (Cell Signaling Technology). The anti-actin antibody was directly conjugated to HRP. The blots were washed in TBS-0.05% tween-20 and incubated in SuperSignal West Pico PLUS HRP Chemiluminescent substrate (Thermo Fisher Scientific) and visualized on a G:BOX by Syngene or a ChemiDoc XRS Imagining system (BioRad). Quantification was done in Image Studio Lite v5.2. Blots were stripped using a guanidine hydrochloride-based (GnHCl) stripping solution (6 M GnHCl, 0.2% Nonidet P-40 (NP-40), 0.1 M β-mercaptoethanol, 20 mM Tris-HCl, pH7.5) (47).

### Statistics

Statistical analyses were performed in GraphPad Prism 10 using the unpaired Student’s t test or a Mann-Whitney *U* test as indicated in the figure legends. P-values <0.05 (*) were considered statistically significant. P-values <0.01 are represented by (**) and P-values <0.001 are represented by (***).

## Results

### Ars/A1 B cells require Lyn to become anergic

To determine if Lyn is required to establish B cell anergy in the Ars/A1 model, we crossed Ars/A1 BCR transgenic mice with mice that have a B cell restricted Lyn deficiency (mb1Cre x Lyn^flox/flox^). Compared to Lyn sufficient littermates (Lyn^flox/flox^ x Ars/A1), B cell intrinsic loss of Lyn in Ars/A1 mice (mb1cre x Lyn^flox/flox^ x Ars/A1) resulted in a drastic reduction of splenic Ars/A1 B cells (Fig 1A). This reduction is not due to aberrant B cell development, since Lyn sufficient and deficient Ars/A1 mice have comparable numbers of immature B cells in their bone marrow (Fig 1B, Suppl Fig 1), while the number of mature recirculating B cells was reduced in Lyn-deficient Ars/A1 mice (Fig 1C). Despite having fewer B cell in the periphery, Lyn deficient Ars/A1 mice have significantly more autoantibody secreting B cells in the spleen compared to Lyn-sufficient littermates, measured both as absolute number of antibody forming cells/spleen (Fig 1D) as well as when normalized to the number of Ars/A1 B cells/spleen (Fig 1E). This suggests that there is a defect in the ability to tolerize Lyn-deficient Ars/A1 B cells. Phenotypically, Lyn deficient Ars/A1 B cells have slightly elevated levels of surface antigen receptors and the activation markers CD69, CD86 and CD95 (Fig 1F). Upon BCR crosslinking, Lyn-deficient Ars/A1 B cells were able to efficiently elevate intracellular calcium concentrations (Fig 1G), consistent with an inability to acquire an anergic state. Further analysis of BCR signaling revealed that some signaling events were significantly enhanced (phosphorylation of Akt and Erk) in Lyn deficient Ars/A1 B cells while other signaling events were minorly affected (phosphorylation of Syk) (Fig 1H). Taken together these data demonstrate that Ars/A1 B cells require Lyn to acquire an anergic state.

**Figure 1.**
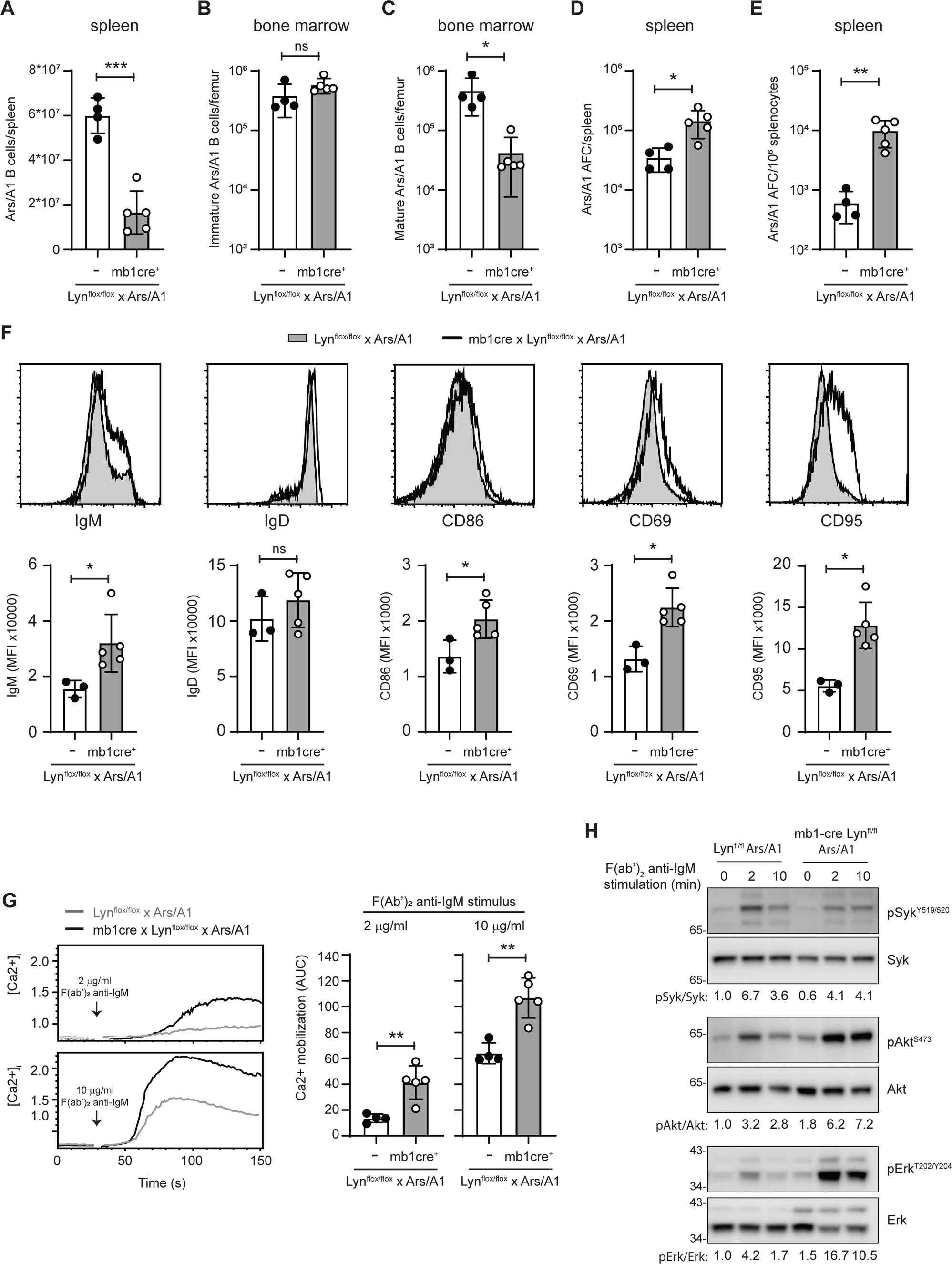
Lyn is required to establish anergy in Ars/A1 B cells. 8-10 wks old mb1Cre x Lyn^flox/flox^ x Ars/A1 and Lyn^flox/flox^ x Ars/A1 littermates were analyzed (n=4-5). The total number of Ars/A1 B cells per spleen were determined (A), as well as the total number of immature Ars/A1 B cells (B) and mature Ars/A1 B cells (C) per femur. The total number of antibody secreting Ars/A1 B cells per spleen (D) and per million splenic Ars/A1 B cells (E) was determined by ELISPOT. F) B cell surface expression of the indicated proteins was determined in a cohort of 8-12 wks old mb1Cre x Lyn^flox/flox^ x Ars/A1 and Lyn^flox/flox^ x Ars/A1 littermates (n=3-5). Representative histograms (gated on B220+) are shown at the top and MFIs are plotted below. G) Calcium mobilization was determined in B cells from a cohort of 8-10 wks old mb1Cre x Lyn^flox/flox^ x Ars/A1 mice and Lyn^flox/flox^ x Ars/A1 littermates (n=4-5) following BCR crosslinking with the indicated doses of anti-IgM F(ab’)_2_. Representative calcium responses (gated on B220+ cells) are shown on the left and the area under the curve is plotted on the right. H) B cells were isolated from mb1Cre x Lyn^flox/flox^ x Ars/A1 mice and Lyn^flox/flox^ x Ars/A1 littermates and stimulated with 10 μg/mL F(ab’)_2_ anti-IgM for the indicated time. Phosphorylation of the indicated proteins was determined by immunoblotting. Following stripping the blots were reprobed for total protein. Data shown are representative of at least three independent replicate experiments. Error bars represent mean ± SD. Two-tailed unpaired Student’s *t* test was used. ns, P > 0.05; *, P < 0.05; **, P < 0.01; ***, P < 0.001.

### Lyn is required for the maintenance of B cell unresponsiveness by suppressing PI3K-dependent signaling events in anergic Ars/A1 B cells

Our analysis of mb1cre x Lyn^flox/flox^ Ars/A1 mice suggests that loss of Lyn does not affect all signaling events downstream of the BCR equally (Fig 1H). To gain more insight into the mechanisms by which Lyn restricts BCR signaling in anergic B cells, we analyzed the impact of Lyn deficiency on BCR signaling in Ars/A1 B cells in more detail. To minimize confounding effects due to a lack of proper tolerization of Ars/A1 B cells during development (Fig 1), and possible adaptation to compensatory mechanisms by other Src family kinases, we used an inducible cre to delete Lyn in mature anergic Ars/A1 B cells. Ars/A1 mice were crossed with mice that express a B cell-restricted inducible cre (hCD20-CreTAM) and have a YFP reporter gene (Rosa26-stopflox YFP) and floxed Lyn locus (Fig 2A, Lyn KO Ars/A1). Seven days following tamoxifen treatment, splenic YFP+ B cells from these mice were Lyn deficient (Fig 2B). For controls we generated Ars/A1 mice that express an inducible Cre and have a YFP reporter and WT Lyn locus. Following tamoxifen treatment, these Ars/A1 B cells express YFP but remain Lyn sufficient (Fig 2A-B, WT Ars/A1). For comparison we also included mice with a WT repertoire of BCRs that upon tamoxifen treatment express YFP but remain Lyn sufficient (Fig 2A, WT).

**Figure 2.**
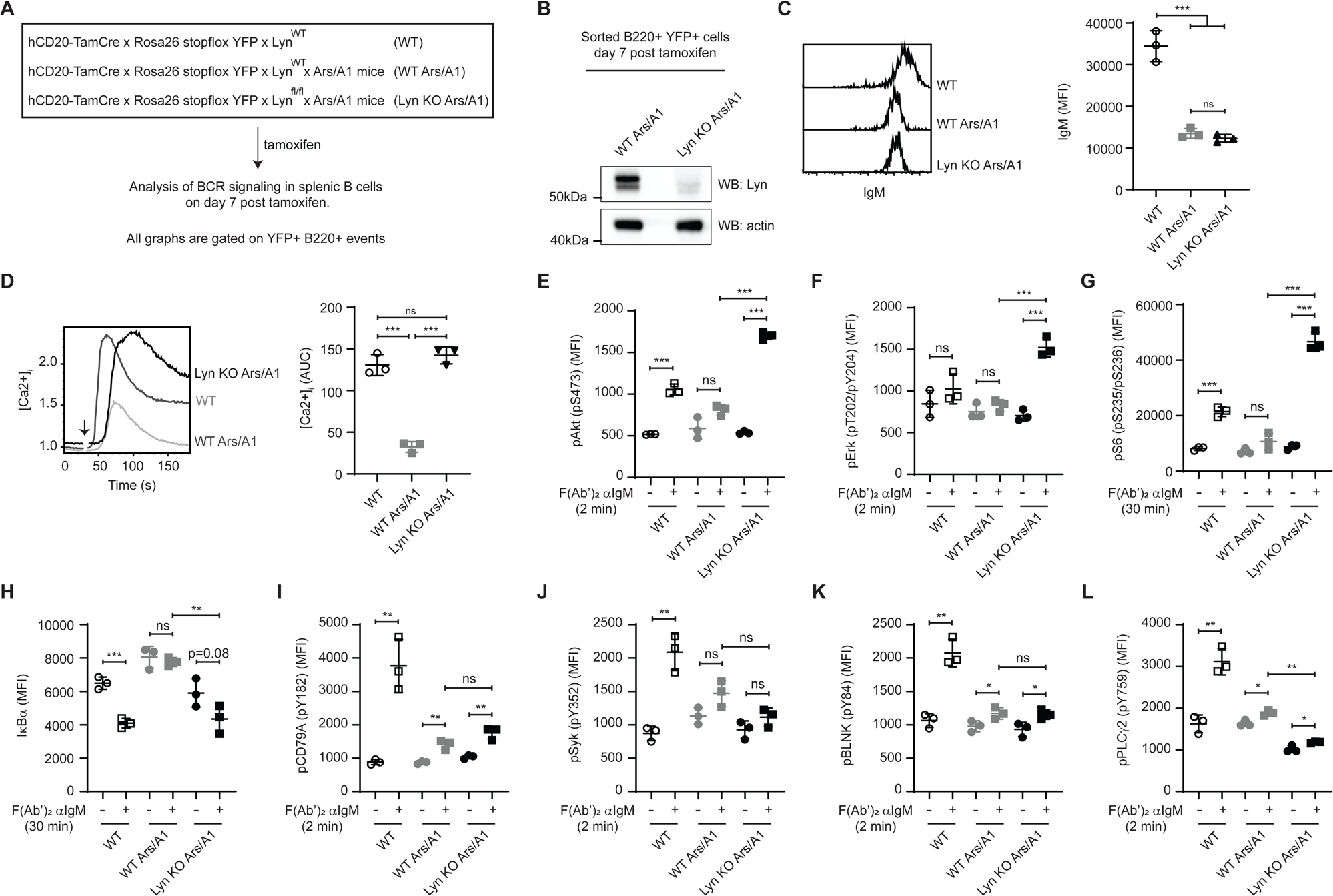
Acute deletion of Lyn restores distal but not receptor proximal signaling events in Ars/A1 B cells. A) Schematic representation of the experimental design. B) YFP+ B cells were sorted seven days post tamoxifen treatment from the indicated mice and Lyn expression was determined by immunoblot. C) Surface IgM expression on B220+ YFP+ cells was determined seven days post tamoxifen treatment (n=3/group). Representative histograms are shown on the left and MFIs are plotted on the right. D) Calcium mobilization was determined in B220+ YFP+ B cells following BCR crosslinking with 5 μg/mL anti-IgM F(ab’)_2_ (n=3/group). Representative calcium responses are shown on the left and the area under the curve is plotted on the right. E-L) Splenocytes were stimulated for the indicated time with 10 μg/mL anti-IgM F(ab’)_2_ and phosphorylation (E-G, I-L) or protein degradation (H) of the specified proteins was measured by intracellular flow cytometry in B220+ YFP+ cells (n=3/group). Data shown are representative of at least two independent replicate experiments. Error bars represent mean ± SD. Two-tailed unpaired Student’s *t* test was used. ns, P > 0.05; *, P < 0.05; **, P < 0.01; ***, P < 0.001

Inducible deletion of Lyn did not significantly alter surface expression of IgM BCRs on Ars/A1 B cells (Fig 2C). Therefore, any observed changes in signaling in these Ars/A1 B cells are due to a loss of Lyn function and not to changes in the degree of BCR crosslinking. In our experiments we used a polyclonal F(ab’)_2_ anti-IgM as a stimulus to avoid any confounding effects due to FcγRIIB recruitment. As observed in Ars/A1 mice that lost Lyn expression early during B cell development (Fig 1G), acute loss of Lyn restores the ability of Ars/A1 B cells to efficiently mobilize calcium following BCR stimulation (Fig 2D). Induced Lyn-deficiency also restored (or enhanced) phosphorylation of Akt (Fig 2E, Suppl Fig 2), S6 (Fig 2G) and Erk (Fig 2F), as well as degradation of IκBα (Fig 2H), in Ars/A1 B cells. In contrast, other signaling events (phosphorylation of Igα ITAM tyrosine 182 (Fig 2I), Syk (Fig 2J), BLNK (Fig 2K) and PLCγ2 (Fig 2L)) remained unchanged when compared to Lyn sufficient Ars/A1 B cells. These events were significantly reduced compared to naïve (WT) B cells (Fig 2I-L). To determine if this selective impact of induced Lyn deficiency on BCR signaling was unique to anergic B cells, we did a similar experiment using mice that have a diverse BCR repertoire. We observed a similar pattern in Lyn deficient B cells where phosphorylation of Akt (Fig 3D, Suppl Fig 3), S6 (Fig 3F) and Erk (Fig 3E), as well as degradation of IκBα (Fig 3G) and calcium mobilization (Fig 3C) are enhanced, albeit with a small delay, while the phosphorylation of Igα (Fig 3H), Syk (Fig 3I), BLNK (Fig 3J) and PLCγ2 (Fig 3K) is reduced and delayed following IgM crosslinking.

**Figure 3.**
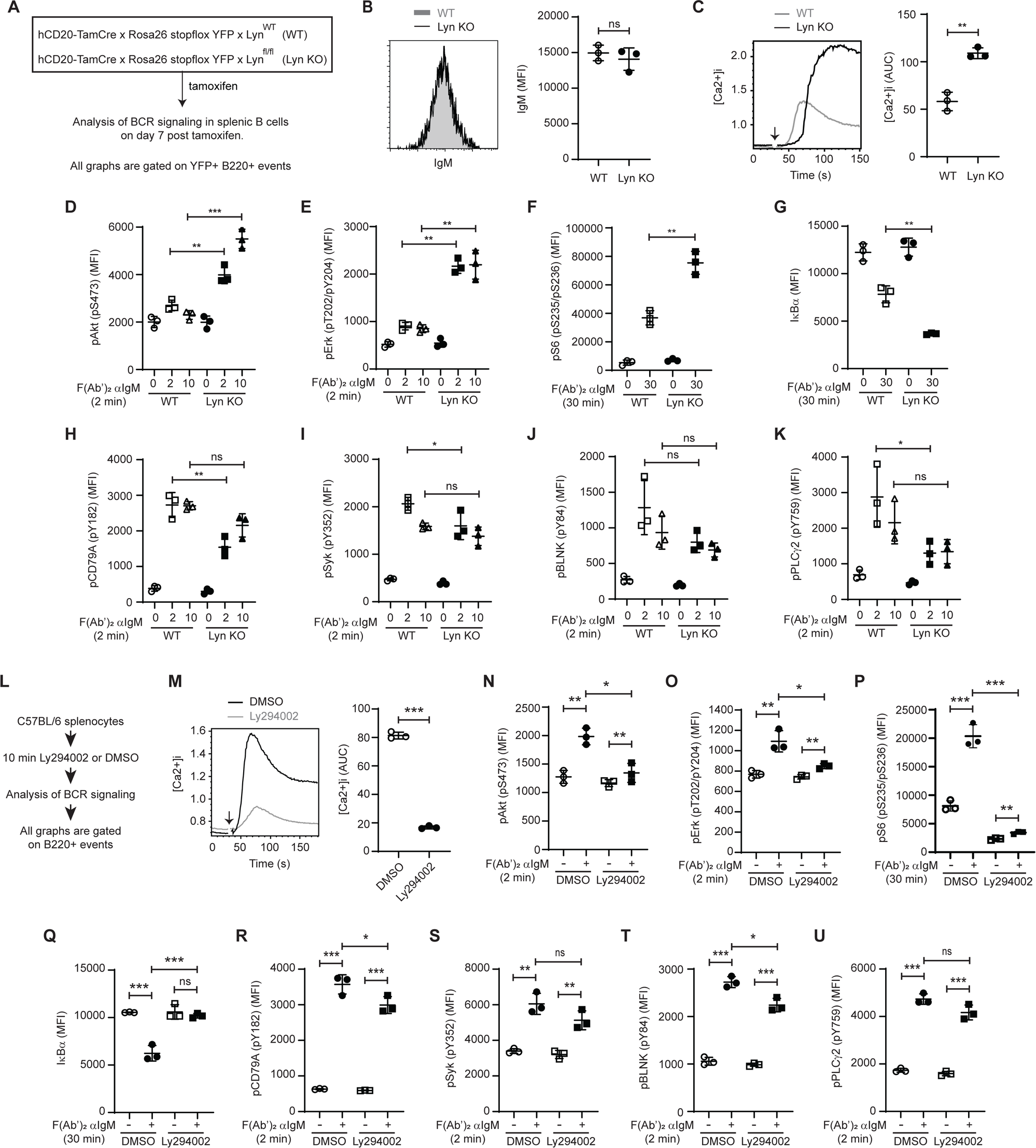
Lyn deficiency primarily enhances PI3K-dependent signaling in B cells. A) Schematic representation of the experimental design. B) Surface IgM expression on B220+ YFP+ cells was determined seven days post tamoxifen treatment (n=3/group). Representative histograms are shown on the left and MFIs are plotted on the right. C) Calcium mobilization was determined in B220+ YFP+ B cells following BCR crosslinking with 5 μg/mL anti-IgM F(ab’)_2_ (n=3/group). Representative calcium responses are shown on the left and the area under the curve is plotted on the right. D-K) Splenocytes were stimulated for the indicated time with 10 μg/mL anti-IgM F(ab’)_2_ and phosphorylation (D-F, H-K) or protein degradation (G) of the specified proteins was measured by intracellular flow cytometry by gating on B220+ YFP+ cells (n=3/group). L) Schematic representation of the experimental design. M) Calcium mobilization was determined in B220+ cells following BCR crosslinking with 5 μg/mL anti-IgM F(ab’)_2_ (n=3/group). Representative calcium responses are shown on the left and the area under the curve is plotted on the right. N-U) Splenocytes were stimulated for the indicated time with 10 μg/mL anti-IgM F(ab’)2 and phosphorylation (N-P, R-U) or protein degradation (Q) of the specified proteins was measured by intracellular flow cytometry by gating on B220+ cells (n=3/group). Data shown are representative of at least two independent replicate experiments. Error bars represent mean ± SD. Two-tailed unpaired Student’s *t* test was used. ns, P > 0.05; *, P < 0.05; **, P < 0.01; ***, P < 0.001

The enhancement of select signaling events suggests that Lyn plays an active role in suppressing these events. Lyn has previously been implicated in regulating PI3K-dependent signaling (9, 48). Suppression of PI3K signaling is important for B cell tolerance (21, 39), including B cell anergy. To determine which of the monitored signaling events are dependent on PI3K signaling, we treated B cells from WT C57BL/6 mice with the PI3K inhibitor Ly294002 and performed a similar analysis of BCR signaling. Interestingly, all the signaling events that were restored in Lyn deficient Ars/A1 B cells were either completely (phosphorylation of Akt (Fig 3N, Suppl Fig 3), S6 (Fig 3P) and degradation of IκBα (Fig 3Q)) or largely (phosphorylation of Erk (Fig 3O) and calcium mobilization (Fig 3M)) dependent on PI3K signaling. The signaling events that were not impacted by Lyn deletion in Ars/A1 B cells (phosphorylation of Igα ITAM tyrosine 182 (Fig 3R), Syk (Fig 3S), BLNK (Fig 3T) and PLCγ2 (Fig 3U)) were not impacted by inhibition of PI3K signaling either. The small significant change in Igα (Fig 3R) and BLNK (Fig 3T) phosphorylation upon BCR crosslinking was not a reproducible finding. Taken together these data show that continuous expression of Lyn is required to maintain BCR unresponsiveness in Ars/A1 B cells and suggests that Lyn primarily operates by regulating PI3K-dependent signaling in anergic Ars/A1 B cells.

### IgM downregulation and suppression of PI3K signaling are responsible for the suppressed BCR signaling phenotype of Ars/A1 B cells

Some signaling events are reduced in Ars/A1 B cells regardless of the presence or absence of Lyn (Fig 2I-L). Acute deletion of Lyn did not alter the surface expression of IgM BCR on Ars/A1 B cells, which remained significantly lower than most B cells within the WT repertoire (Fig 2C). Downregulation of surface IgM BCR is one of the ways by which autoreactive B cells are thought to reduce antigen receptor signaling strength to stay below an activation threshold.

To determine how differences in IgM expression contribute to the difference in signaling between WT repertoire B cells and Ars/A1 B cells, we used a protocol that allows us to compare subpopulations of B cells that have equal levels of surface IgM BCR and therefore equal BCR crosslinking. Prior to stimulation with a polyclonal F(ab’)_2_ anti-IgM, we stained B cells with a non-stimulatory fluorescently tagged Fab fragment of an anti-IgM monoclonal antibody, clone b76 (Fig 4A). Analysis of the phosphorylation of the Y182 ITAM residue of CD79A shows that at a fixed dose of crosslinking antibody there is a direct correlation between the amount of IgM BCR expressed on a B cell and the magnitude of induced phosphorylation (Suppl Fig 4A). BCR internalization following BCR crosslinking did not affect the detection of fluorescently tagged Fab anti-IgM during the first 45 minutes following BCR crosslinking (Suppl Fig 4B). Therefore, we assume in our analysis that the Fab anti-IgM signal measured following stimulation represents the IgM surface expression levels at the time of stimulation.

**Figure 4.**
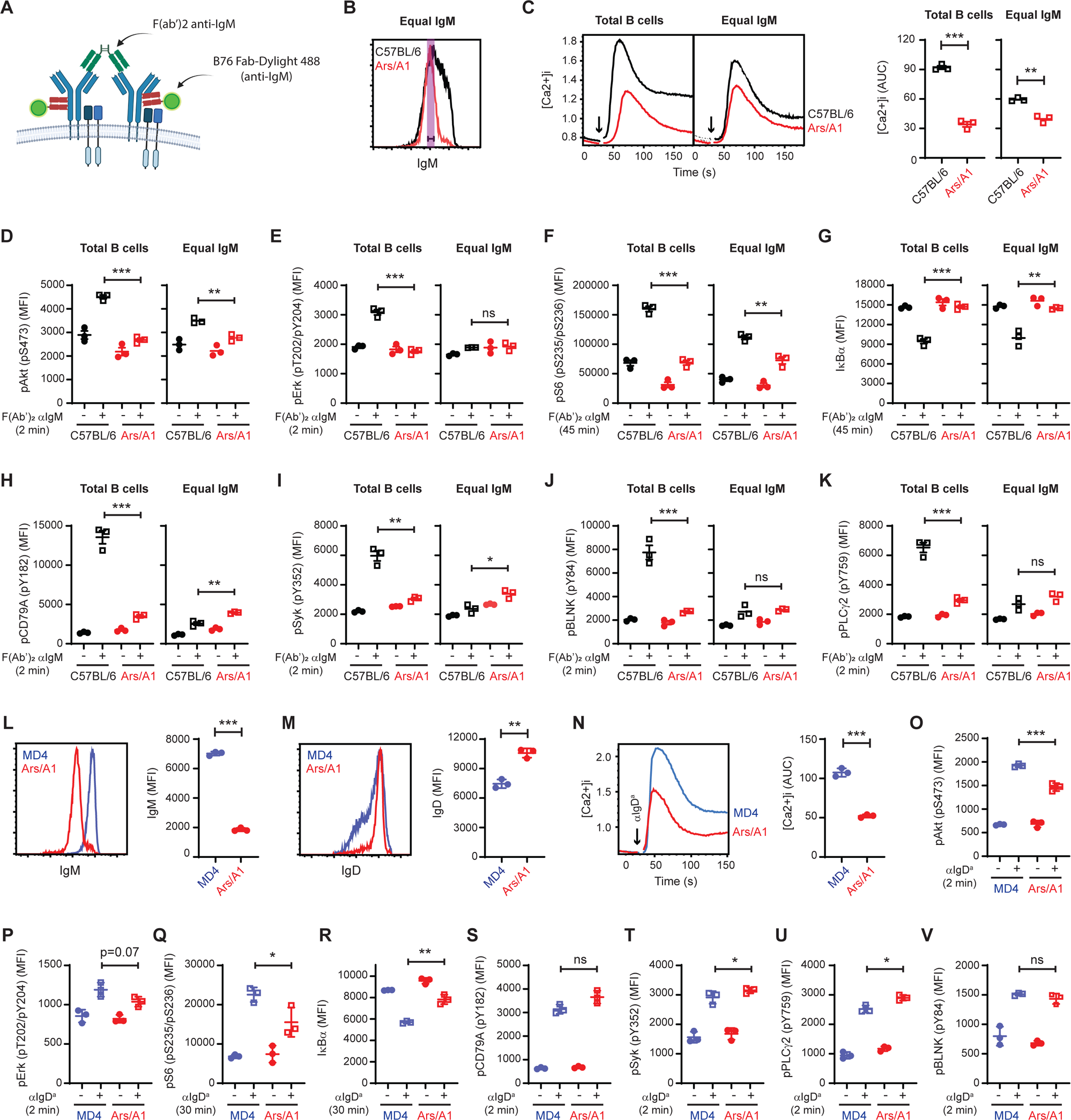
Reduced IgM expression and active suppression of PI3K dependent signaling contribute to the suppressed signaling phenotype of anergic B cells. A) Detection of IgM BCR expression with a non-stimulatory Fab anti-IgM-Dylight488 fragment. B) Surface IgM expression on B220+ C57BL/6 (black) and Ars/A1 B cells (red). Gating for comparison of signaling in equal IgM expressing B cells is indicated. C) Calcium mobilization following BCR crosslinking with 5 μg/mL anti-IgM F(ab’)_2_. Representative calcium responses are shown on the left, gated on the total B cell population (B220+) and on B cells expressing equal IgM, the area under the curve is plotted on the right (n=3). D-K) Splenocytes were stimulated for the indicated time with 10 μg/mL anti-IgM F(ab’)_2_ and phosphorylation (D-F, H-K) or protein degradation (G) of the specified proteins was measured by intracellular flow cytometry. MFIs are plotted of the total B cell population (B220+) and of B cell expressing equal IgM (n=3/group). L-V) Signaling in MD4 (blue) vs Ars/A1 (red) B cells following IgD stimulation. Surface IgM (L) and IgD (M) on B220+ MD4 and Ars/A1 B cells. Representative plots are shown on the left and MFIs are plotted on the right. N) Calcium mobilization was determined in B220+ cells following BCR crosslinking with 5 μg/mL anti-IgD^a^ (n=3/group). Representative calcium responses are shown on the left and the area under the curve is plotted on the right. O-V) Splenocytes were stimulated for the indicated time with 5 μg/mL anti-IgD^a^ and phosphorylation (O-Q, S-V) or protein degradation (R) of the indicated proteins was measured by intracellular flow cytometry by gating on B220+ cells (n=3/group). Data shown are representative of at least two independent replicate experiments. Error bars represent mean ± SD. Two-tailed unpaired Student’s *t* test was used. ns, P > 0.05; *, P < 0.05; **, P < 0.01; ***, P < 0.001

As reported previously (41), when we compared BCR signaling in the <u>total</u> population of B cells, all signaling events monitored were significantly reduced in Ars/A1 B cells compared to naïve WT C57BL/6 B cells (Fig 4C-K, total B cell plots). However, when we analyze subpopulations with equal IgM BCR expression, differences emerge. Some signaling events remain significantly lower in Ars/A1 B cells such as Akt phosphorylation (Fig 4D, Suppl Fig 4), calcium mobilization (Fig 4C) and IκBα degradation (Fig 4G). In contrast, other events such as BLNK (Fig 4J) or PLCγ2 (Fig 4K) phosphorylation were comparable between Ars/A1 B cells and WT C57BL/6 B cells expressing equal surface IgM, and phosphorylation of Igα (Fig 4H) and Syk (Fig 4I) are even slightly increased in Ars/A1 B cells. These results suggest that downregulation of surface IgM BCR accounts for much of the impaired BCR signaling phenotype we observe in Ars/A1 B cells, while additional active mechanisms suppress PI3K dependent signaling further.

One caveat of this analysis is that the subpopulation of B cells within the WT repertoire that express low IgM are enriched for autoreactive B cells (49). These autoreactive B cells may have their own active suppression mechanisms that could mask the detection of inhibitory mechanisms in Ars/A1 B cells. To circumvent this potential confounding factor, we did a parallel experiment using B cells from MD4 mice. MD4 mice express a transgenic BCR that binds Hen Egg Lysozyme (HEL) with a high affinity and has little cross reactivity with self-antigen. MD4 B cells express very high surface IgM levels (Fig 4L), with virtually no overlap with Ars/A1 B cells, making it unfeasible to compare subpopulation with equal IgM expression. However, IgD BCR expression levels are more comparable between MD4 and Ars/A1 B cells (Fig 4M), with Ars/A1 B cells expressing slightly higher IgD levels. When we analyzed signaling induced following IgD crosslinking, we found that while most signaling events were comparable between MD4 and Ars/A1 B cells (Fig 4S-V, Suppl. Fig 4), PI3K dependent events were significantly suppressed (Fig 4N-R). Taken together these results suggest that downregulation of IgM BCR levels contributes significantly to the suppressed BCR signaling phenotype of anergic Ars/A1 B cells, but that active suppression of PI3K signaling via Lyn also plays an important role.

### Acute deletion of Lyn in Ars/A1 B cells results in inefficient autoantibody responses

Acute deletion of Lyn in anergic Ars/A1 B cells restores their ability to signal following BCR stimulation (Fig 2). Previously, we used an adoptive transfer approach (40) to determine if SHP-1 and SHIP-1 were required to maintain B cell anergy and showed that following acute deletion of either of these phosphatases, Ars/A1 B cells mounted an autoantibody response. Using a similar adoptive transfer approach (Fig 5A), we observed that following acute Lyn deletion there was a significant reduction in the number of YFP+ (Lyn-deficient) Ars/A1 B cells (suppl Fig 5A-B and Fig 5D) in the spleen over time. At day 14 post tamoxifen treatment the remaining YFP+ Lyn-deficient Ars/A1 B cells had a mixed phenotype. In some of the recipient mice few YFP+ Lyn KO Ars/A1 B cells had proliferated, while in other mice the majority had proliferated and a fraction of those had differentiated into plasmablasts. The absolute numbers of cells were so low that in most mice it did not result in a detectable antibody secreting cell response. We did 4 replicate experiments in which we adoptively transferred variable numbers of Ars/A1 B cells (Suppl Fig 5C-F). In all experiments we observed a considerable variation in the response between mice within each group. There was a trend towards more pronounced autoantibody responses when more Ars/A1 B cells were transferred (Suppl Fig 5), but the overall cell recovery remains low. To exclude the possibility that there is an inherent problem in mounting an antibody response following acute Lyn deletion, we compared the ability of Lyn sufficient or deficient MD4 B cells to respond to SRBC-HEL immunization (Fig 5F-J). To be able to directly compare antibody secreting cell responses we sorted YFP+ MD4 B cells that were Lyn sufficient or deficient and adoptively transferred equal numbers. Following immunization, we observed that Lyn deficient MD4 B cells have a higher frequency of plasmablasts (Fig 5H, J) but overall mounted an equal antibody forming cell response (Fig 5G). These results suggest that while acute loss of Lyn results in a loss of B cell anergy, the resultant autoantibody responses are inefficient, possibly due to reduced fitness of B cells. Finally, we determined if induced Lyn haploinsufficiency promotes autoantibody responses by itself or when combined with induced haploinsufficiency of SHIP-1 (Fig 5K-O). While induced haploinsufficiency of either Lyn or SHIP-1 in isolation did not induce significant proliferation of Ars/A1 B cells or accumulation of autoantibody secreting cells, induced double haploinsufficiency of Lyn and SHIP-1 in Ars/A1 B cells did (Fig 5O). Under these conditions no significant loss of YFP+ Ars/A1 B cells was observed (Fig 5N). Collectively these data suggest that Lyn function is required for the maintenance of B cell anergy and that reductions in Lyn function can synergize with reductions of other proteins in the pathway to result in a loss of tolerance of anergic B cells.

**Figure 5.**
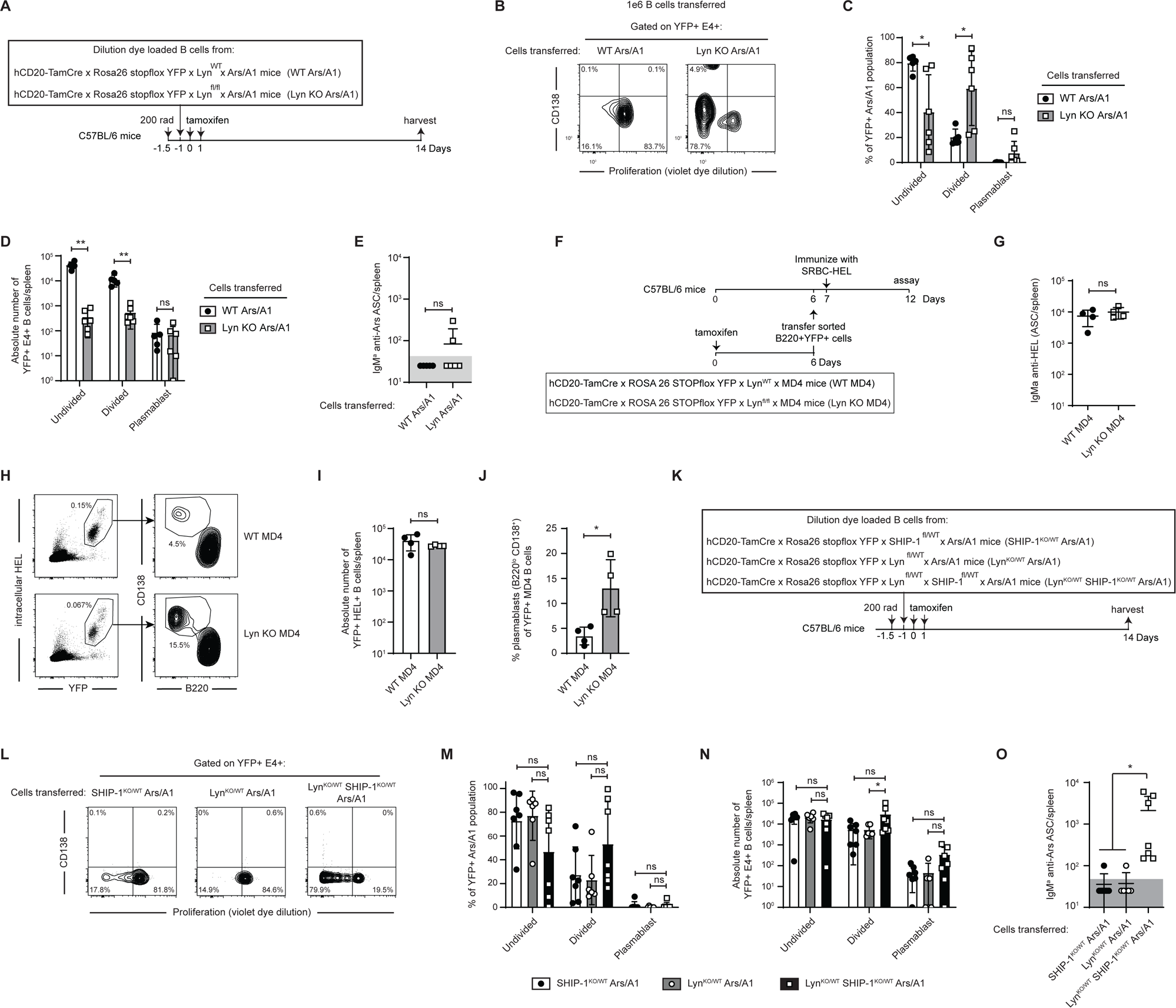
Acute deletion of Lyn results in inefficient autoantibody responses. A) Schematic representation of the experimental protocol. B) Proliferation and differentiation to plasmablasts of splenic YFP+ E4+ Lyn sufficient (WT Ars/A1) and Lyn deficient (Lyn KO Ars/A1) B cells 14 days after tamoxifen treatment (n=5-6/group, representative cytograms shown). C) Distribution of the YFP+ E4+ B cell population shown in Fig 5B between an undivided, proliferated and plasmablast (proliferated and CD138+) state. D) The absolute number of YFP+ E4+ B cell cells per spleen from Fig 5B in an undivided, proliferated and plasmablast (proliferated and CD138+) state 14 days after tamoxifen treatment. E) Quantification of antibody secreting cells (IgM^a^ anti-Ars) by ELISPOT of spleen cells from Fig 5B. Gray area delineates the limit of detection (50 spots/spleen). F) Schematic representation of the experimental protocol. G-J) 5 days following immunization with SRBC-HEL spleens (n=4) were analyzed for G) number of antibody secreting cells (IgM^a^ anti-HEL) by ELISPOT, H) the frequency of YFP+ MD4 B cells and plasmablasts (B220^lo^ CD138^+^), I) the absolute number of YFP+ MD4 B cells per spleen and J) the frequency of plasmablasts of the YFP+ MD4 population. K) Schematic representation of the experimental protocol. L) Proliferation and differentiation to plasmablasts of splenic YFP+ E4+ SHIP-1 haploinsufficient (SHIP-1^KO/WT^ Ars/A1), Lyn haploinsufficient (Lyn^KO/WT^ Ars/A1) and Lyn and SHIP-1 double haploinsufficient (SHIP-1^KO/WT^ Lyn^KO/WT^ Ars/A1) B cells 14 days after tamoxifen treatment (n=6-7/group, representative cytograms shown). M) Distribution of the YFP+ E4+ B cell population shown in Fig 5L between an undivided, proliferated and plasmablast (proliferated and CD138+) state. N) The absolute number of YFP+ E4+ B cell cells per spleen from Fig 5L in an undivided, proliferated and plasmablast (proliferated and CD138+) state 14 days after tamoxifen treatment. O) Quantification of antibody secreting cells (IgM^a^ anti-Ars) by ELISPOT of spleen cells from Fig 5L. Gray area delineates the limit of detection (50 spots/spleen). Data shown are representative of at least two replicate experiments. Error bars represent mean ± SD. Two-tailed unpaired Student’s *t* test was used in Fig 5C, J and M). A Mann-Whitney *U* test was used in Fig 5D, E, I, N and O. ns, P > 0.05; *, P < 0.05; **, P < 0.01; ***, P < 0.001.

## Discussion

Lyn deficient mice develop autoantibodies and lupus-like disease (8, 12, 13). Multiple cell types contribute to this loss of tolerance, including dendritic cells (45) and B cells (11). Importantly, B cell-restricted loss of Lyn is sufficient to result in a loss of B cell tolerance and associated autoantibody deposition in tissue (11). It is still unclear which B cell tolerance mechanisms are dependent on Lyn function. There is some evidence that central tolerance mechanisms are impacted by Lyn dysregulation, resulting in a higher frequency of autoreactive B cells in the periphery (50, 51). Among the peripheral B cell populations that harbor autoreactive B cells both B-1a (52, 53) and T-bet expressing Age-associated B Cells (ABC-B cells) (54) contribute to the autoantibody producing cell population in Lyn-deficient animals.

Starting at the transitional B cell stage (11, 55), Lyn becomes critical for establishing inhibitory signaling mechanisms in B cells. This includes the T3 population which contains anergic autoreactive B cells (32, 56). These inhibitory signaling mechanisms are important for peripheral B cell tolerance (19–21), including B cell anergy (21, 40, 57), a major peripheral B cell tolerance mechanism. One would expect that Lyn deficiency impacts the ability of autoreactive B cells that are normally silenced by anergy to become tolerized. Surprisingly, two previous studies found that B cell tolerance (defined as a lack of autoantibody production) was preserved when two independent models of B cell anergy were crossed onto a total Lyn KO background (30, 31). In VH3H9 heavy chain transgenic mice on the Balb/c background, λ1-expressing B cells are DNA-reactive and anergic (58, 59). Lyn deficiency did not result in a loss of tolerance of λ1-expressing B cells (31) as measured by autoantibody production, although they did regain responsiveness to LPS. Lyn deficiency in the MD4 x ML5 model of B cell anergy (44) resulted in increased peripheral deletion, leaving few mature Lyn deficient MD4 x ML5 B cells left in the periphery to analyze (30). Nevertheless, the authors found no evidence that the remaining Lyn deficient MD4 x ML5 B cells differentiated into autoantibody producing cells, suggesting that immune tolerance was intact (30).

Here we show that Lyn is required to establish and maintain an anergic state in the DNA-reactive Ars/A1 model of B cell anergy. Two differences in experimental design may contribute to this seemingly opposed finding. While the previous studies used conventional Lyn KO to determine the dependence of B cell anergy on Lyn, we used two different Cre lines to delete Lyn in a B cell restricted manner during B cell development (Fig 1) or in mature anergic B cells (Fig 2, 5). Conventional Lyn deficient animals develop a severe inflammatory environment (13), due to dysregulation of macrophages and dendritic cells (45), that may indirectly impact the B cell compartment. Limiting deletion of Lyn to just B-cells, thus avoiding all the myeloid cell activation, may be key to revealing its role in B-cell anergy. Secondly, it has become increasingly clear that B cell anergy exists on a spectrum (37, 60, 61). Presumably individual autoreactive B cell clones can utilize multiple mechanisms to different degrees to acquire a sufficiently high activation threshold to prevent their activation. Ars/A1 B cells represent a subset of anergic B cells that are strongly dependent on inhibitory signaling and may therefore be more sensitive to Lyn deletion than other models tested. Our results demonstrate that anergic B cells are dependent on Lyn to establish and maintain an anergic state.

Analysis of proximal and distal BCR signaling revealed that Lyn is primarily responsible for suppressing PI3K-dependent signaling in anergic B cells (Figs 2, 3). Suppression of PI3K-dependent signaling has emerged as an important mechanism in peripheral B cell tolerance (39). B cell anergy models (21, 39, 57) and (anergic) autoreactive B cells in the wild type B cell repertoire (62, 63) have increased inositol phosphatase activity and dampened PI3K-dependent signaling. This suppressed state is lost in autoreactive B cells of individuals that develop autoimmune disease (63, 64)

Several mechanisms have been proposed by which Lyn can regulate PI3K-dependent signaling. In a negative feedback mechanism, Lyn is reported to constrain Src family kinase activity and thereby PI3K activation (9). Lyn-mediated activation of SHP-1 can dampen PI3K activation by actively suppressing upstream events (65, 66). Finally, Lyn-mediated activation of SHIP-1 restricts PI3K dependent signaling by dephosphorylating the PI3K product PI(3,4,5)P3, dampening BCR signaling (67). Ars/A1 B cells have increased SHIP-1 phosphorylation, which is Lyn-dependent (21). Our genetic complementation experiments (Fig 5K-O) support an important role for the Lyn-SHIP-1 pathway in controlling B cell tolerance/anergy, similar to previous reports (68). How SHIP-1 is activated in anergic B cells is unclear. SHIP-1 is activated following BCR stimulation independently of FcγRIIB (69, 70), a known SHIP-1 activator. It seems that the BCR complex can directly recruit SHIP-1 (71, 72), possibly in a Dok-3/Grb-2 dependent manner (73–75). This raises the possibility that continuous stimulation of the BCR in autoreactive B cells may establish a Lyn-SHIP-1 dependent inhibitory feedback mechanism, independent of additional inhibitory receptors, that is important for maintaining B cell unresponsiveness.

We also observed that IgM downregulation accounted for much of the hyporesponsive signaling phenotype of Ars/A1 B cells (Fig 4). This is consistent with previous reports that suggest IgM downregulation is an effective mechanism to raise the activation threshold of autoreactive B cells (37, 38, 49). However, Lyn-dependent suppression of PI3K signaling proved critical to maintaining a state of unresponsiveness (Figs 3, 4). Following acute Lyn-deletion, IgM levels remained low, and while BCR proximal events were reduced, BCR distal signaling was restored and enhanced (Fig 2). Phenotypically, this resembles recent observations made while studying BCR signaling following engagement with antigen on virus-like particles (76). In this study they found that there was strong signal amplification of distal PI3K-dependent signaling events, due to the ability of such antigens to circumvent Lyn-dependent regulatory signaling mechanisms. Such antigens were able to induce BCR signaling in anergic MD4 x ML5 B cells (76) and induce autoantibody responses (77). This stresses the importance of regulating PI3K dependent signaling to prevent autoantibody responses (78).

Suppression of PI3K-dependent signaling may also play a role in controlling IgD signaling. Anergic B cells express normal levels of IgD BCR, which has the same self-antigen-specificity as IgM BCR. Studies that used mice that only express one subclass of BCR suggest that self-antigens preferentially engage with IgM BCRs (79, 80), possibly due to structural differences between surface IgM and IgD (81). While expression of IgD attenuates the transcriptional program of anergic B cells (82), it may primarily enable anergic B cells to respond to cross-reacting foreign antigen, a process coined clonal redemption (83). Suppression of PI3K signaling, including following IgD BCR stimulation, may help facilitate this careful balance between allowing an autoreactive anergic B cell to participate in an immune response, while ensuring that it is only allowed to differentiate into an antibody secreting cell when the B cell has mutated sufficiently away from its self-reactivity.

B cell restricted deletion of Lyn resulted in an accumulation of autoantibody producing cells (Fig 1). Yet, our experiments in which a finite amount of B cells were adoptively transferred, followed by acute Lyn deletion, suggest that this process is inefficient. Previous studies describe decreased survival of peripheral B cells in Lyn deficient mice (30, 48, 84, 85), in part due to Lyn’s role in negatively regulating the pro-apoptotic protein Bim (86). This would be expected to decrease the likelihood of autoantibody responses by reducing the precursor frequency of B cells that can participate in a response and limit the success rate of activated autoreactive B cells to counteract Bim-dependent activation induced cell death (87). This may disproportionally affect autoreactive B cells. Lyn deficient mice mount relatively normal antibody responses (12, 13, 88) with an increased plasma blast/cell accumulation (89, 90) (Fig 5G-J). Why Lyn-deficient Ars/A1 B cells respond differently than MD4 B cells is unclear. Ars/A1 B cells may be in a more fragile state, coming out of anergy, and their response is dependent on TLR signals (91) and T cell help (92). Lyn is involved in both TLR (45, 93) and CD40 signaling (94, 95). Changes in these pathways would be expected to affect the survival of DNA reactive B cells (96).

Dysregulation of Lyn (22–25) and its regulators (26–29) in B cells is strongly associated with disruptions in immune tolerance and the development of autoimmune disease. While it is clear that dysregulation of Lyn can impact many processes important to immune homeostasis and immune tolerance, our work suggests that it will also impact the ability of a subset of anergic B cells to establish and maintain an unresponsive state, which may result in their participation in the development of autoimmune disease (97–100)

## Acknowledgements

The authors wish to thank Dr. John Cambier for his support during the early stages of this project and for valuable discussion and editing, Dr Clifford Lowell for Lyn^flox/flox^ mice, and Dr. Tinalyn Kupfer, the CU | AMC ImmunoMicro Flow Cytometry Shared Resource, RRID:SCR_021321, and the Cancer Flow Core (Cancer Center Support Grant (P30CA046934)) for technical assistance. This work was supported by National Institutes of Health grants R21AI149019 and R01AI124487, and the University of Colorado Human Immunology and Immunotherapy Initiative.

