## Supplemental Figures 1-5 for "The Src-family kinase Lyn plays a critical role in establishing and maintaining B cell anergy by suppressing PI3K-dependent signaling"

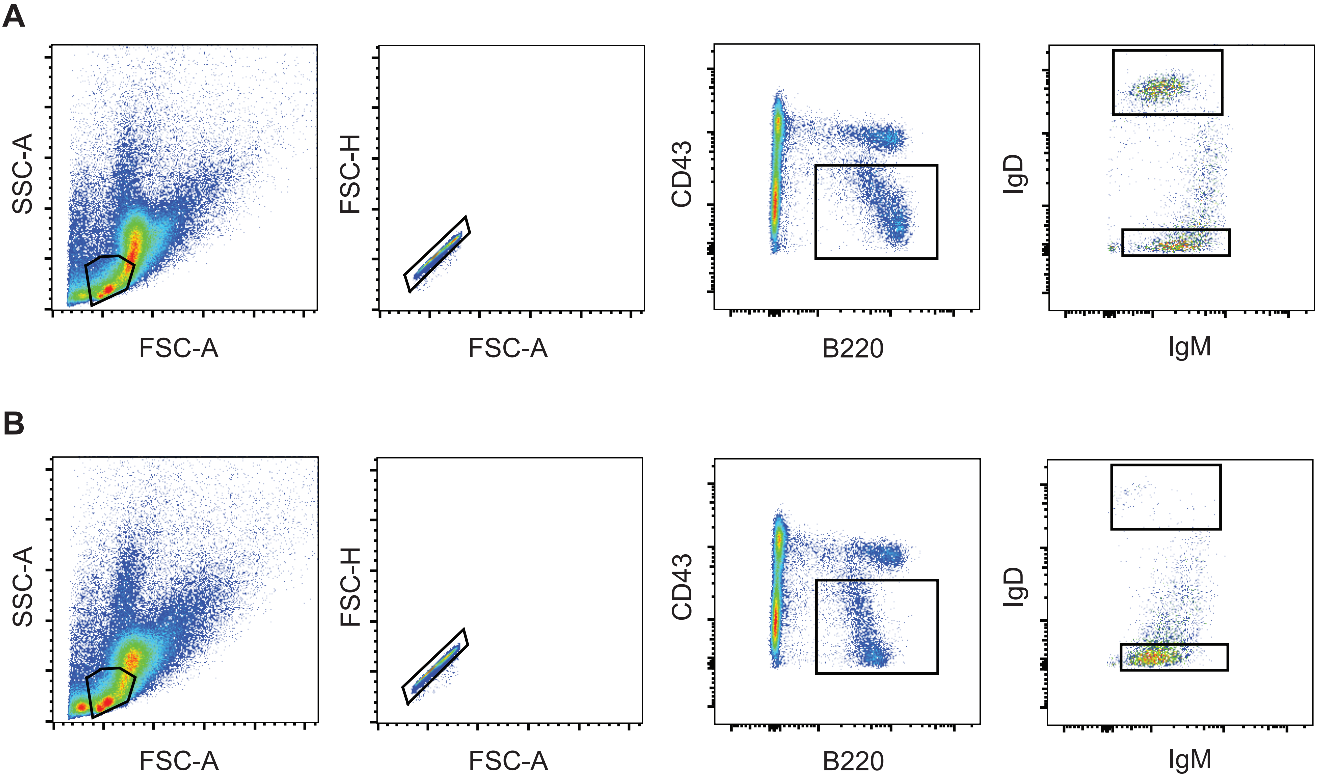


**Supplemental Figure 1. Bone marrow staining gating strategy.** Gating strategy to determine the frequency of immature (B220+ CD43- IgM+ IgD-) and mature (B220+ CD43- IgM+ IgD+) B cells in the bone marrow of Lyn$^{flox/flox}$ x Ars/A1 (A) and mb1cre x Lyn$^{flox/flox}$ x Ars/A1 (B) mice. These frequencies were used to calculate the absolute cell numbers shown in Fig 1B-C. Representative plots are shown.

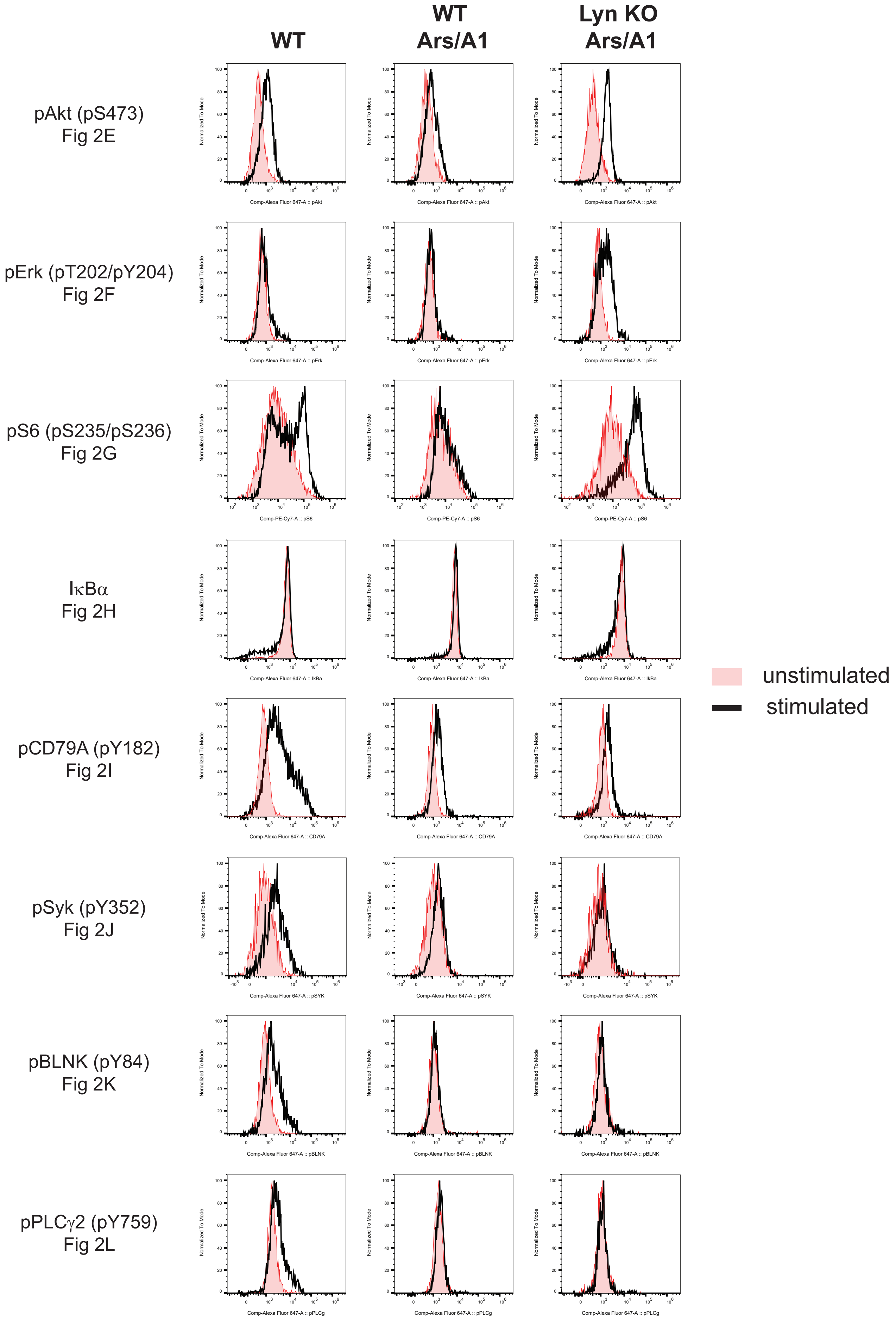


**Supplemental Figure 2. Representative histograms from Fig 2.** Individual histograms were gated on B220+ YFP+ cells (n=3/group). Representative histograms are shown

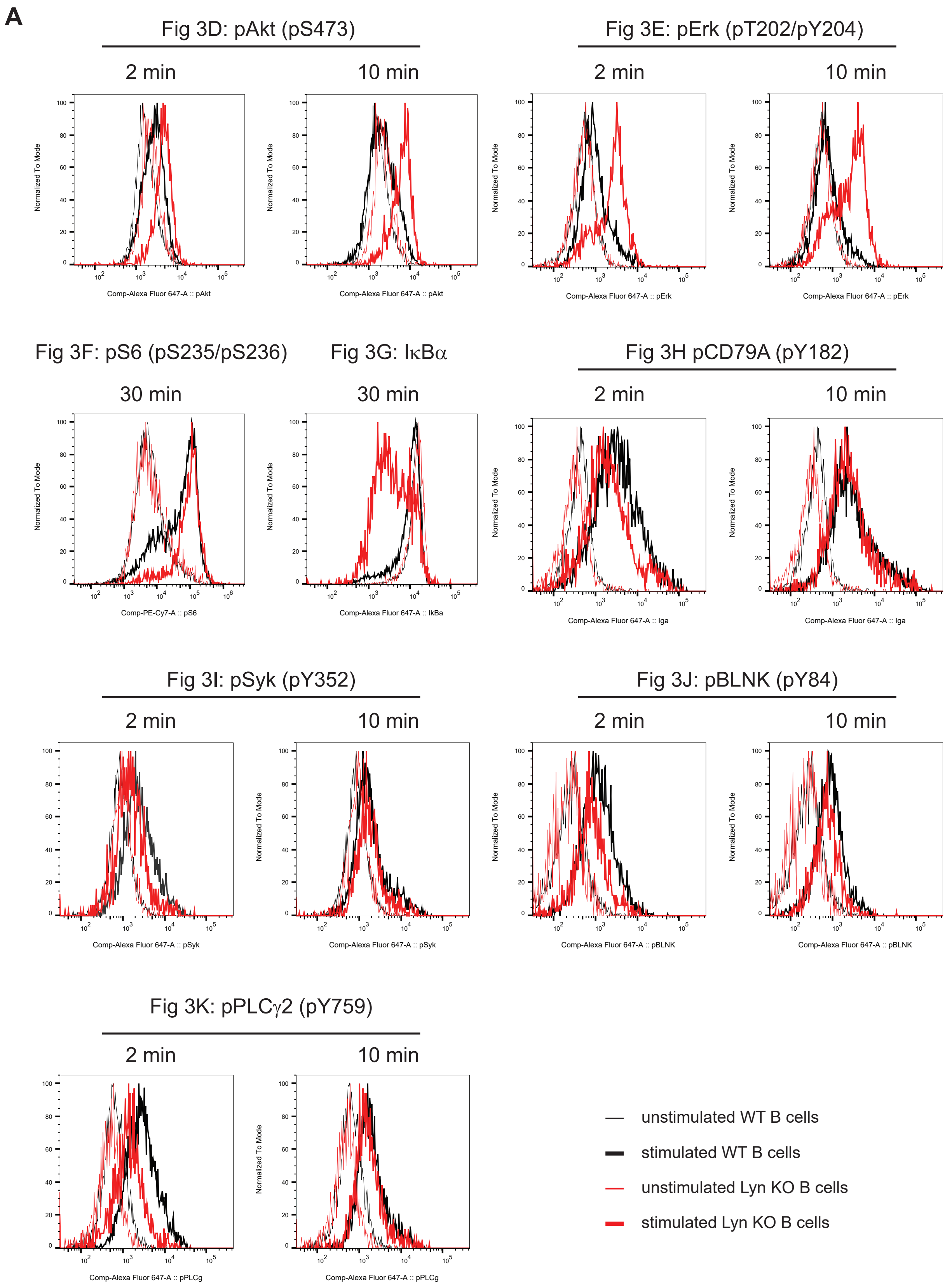


**Supplemental Figure 3A. Representative histograms from Fig 3D-K.** Individual histograms were gated on (A) B220+ YFP+ cells or B) B220+ cells (n=3/group). Representative histograms are shown.

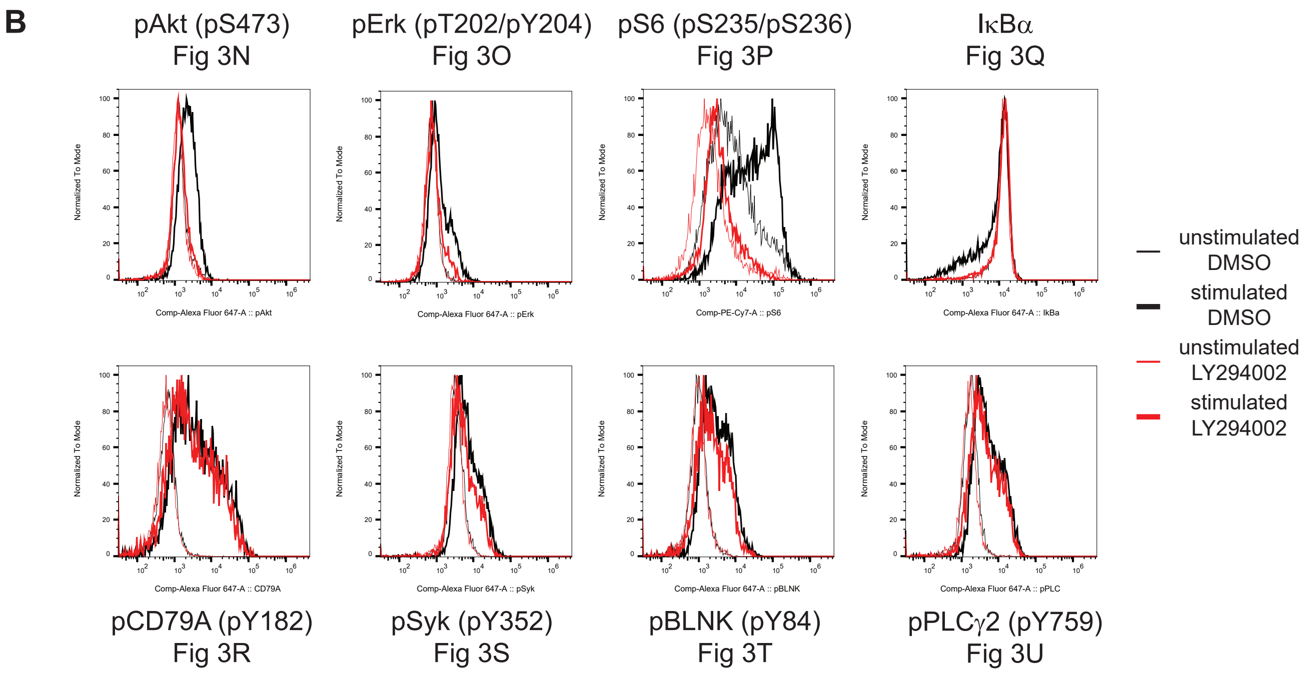


**Supplemental Figure 3B. Representative histograms from Fig 3N-U.** Individual histograms were gated on B220+ cells (n=3/group). Representative histograms are shown

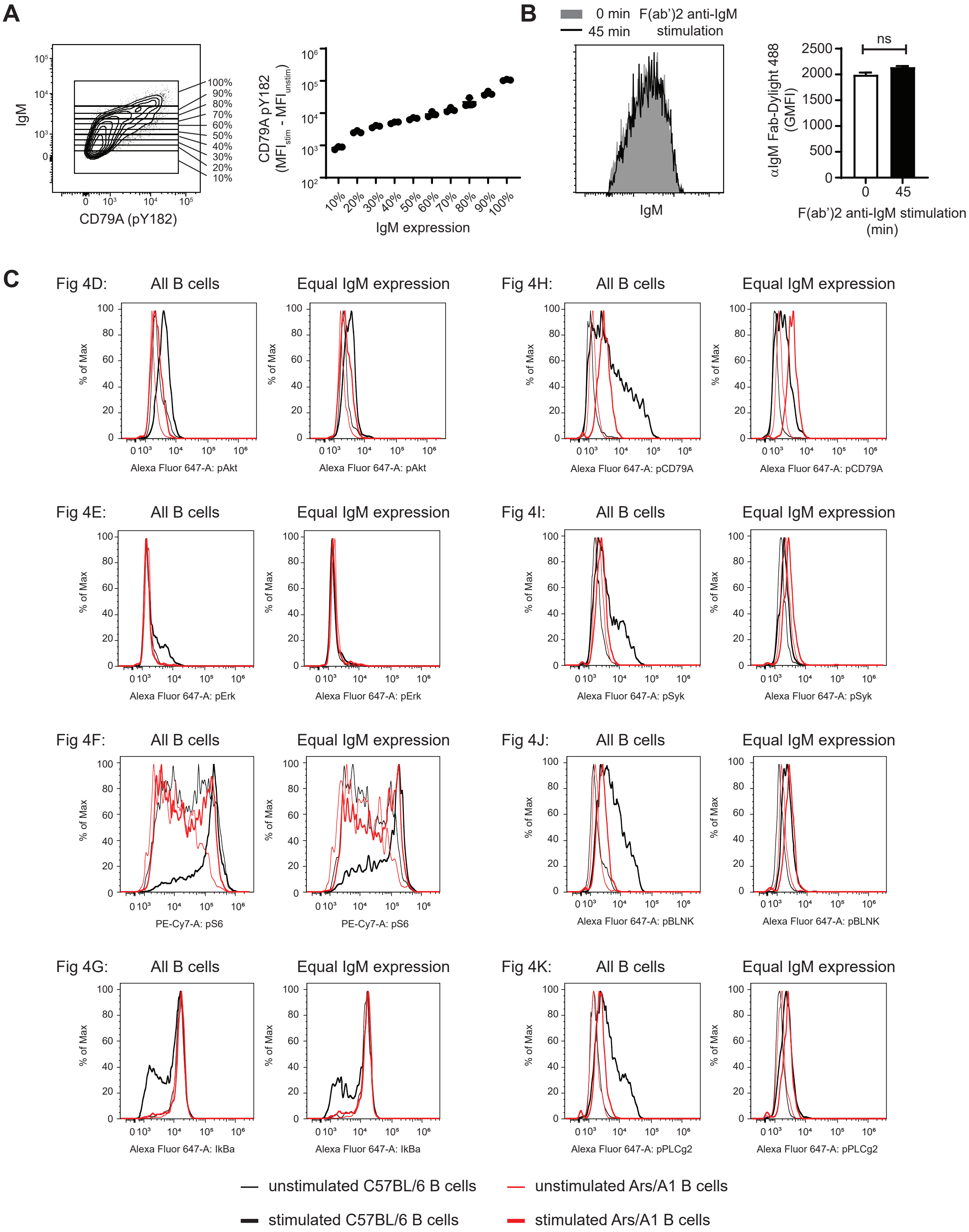


**Supplemental Figure 4A-C.** A) C57BL/6 B cells were stained with anti-IgM Fab fragment-Dylight488 prior to stimulation with $F(ab')_2$ anti-IgM for two minutes. CD79A Y182 phosphorylation was measured by flow cytometry. A representative plot is shown on the left (gated on B220+). The population was divided into 10 equal populations. The increase in MFI is plotted to the right (n=3). B) C57BL/6 B cells were stained with anti-IgM Fab fragment-Dylight488 prior to stimulation with $F(ab')_2$ anti-IgM for 45 minutes. IgM staining at 45 minutes was compared to unstimulated B cells. Representative plot shown to the left and GMFI are plotted on the right. Unpaired t-test was used, $p>0.05$.
C) representative histograms from Fig 4D-K. Individual histograms were gated on B220+ cells.

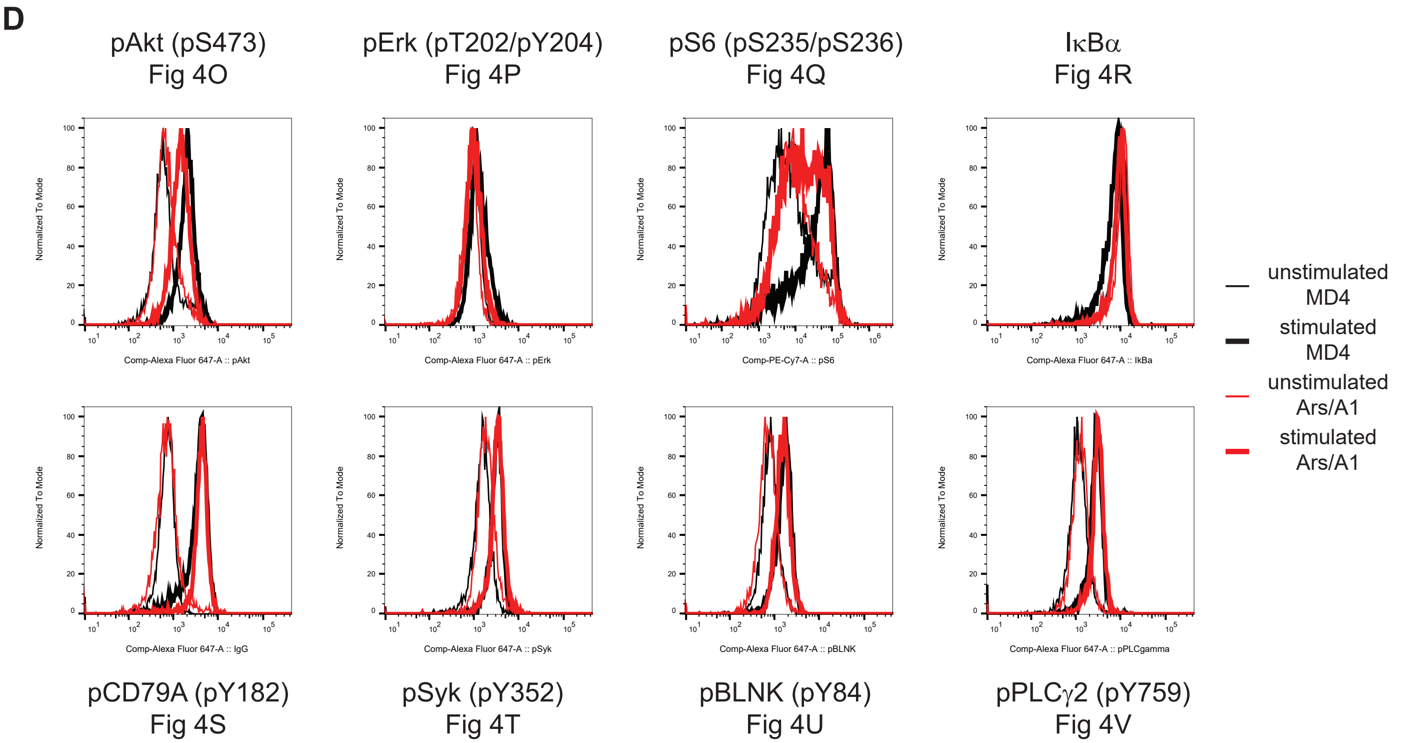


**Supplemental Figure 4D** D) Representative histograms from Fig 4O-V. Individual histograms were gated on B220+cells (n=3/group).

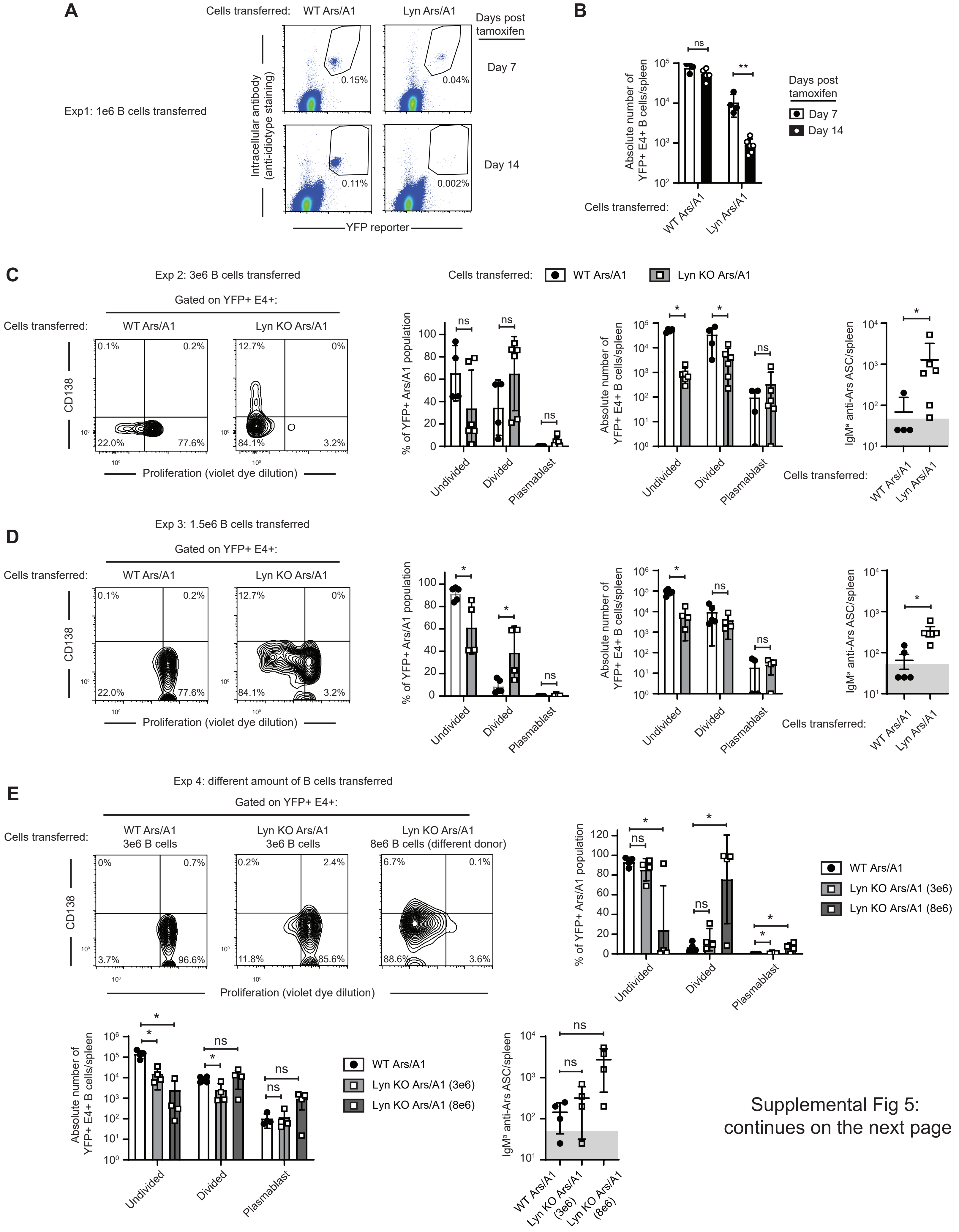


Supplemental Fig 5:
continues on the next page

Supplemental Fig 5 continued.

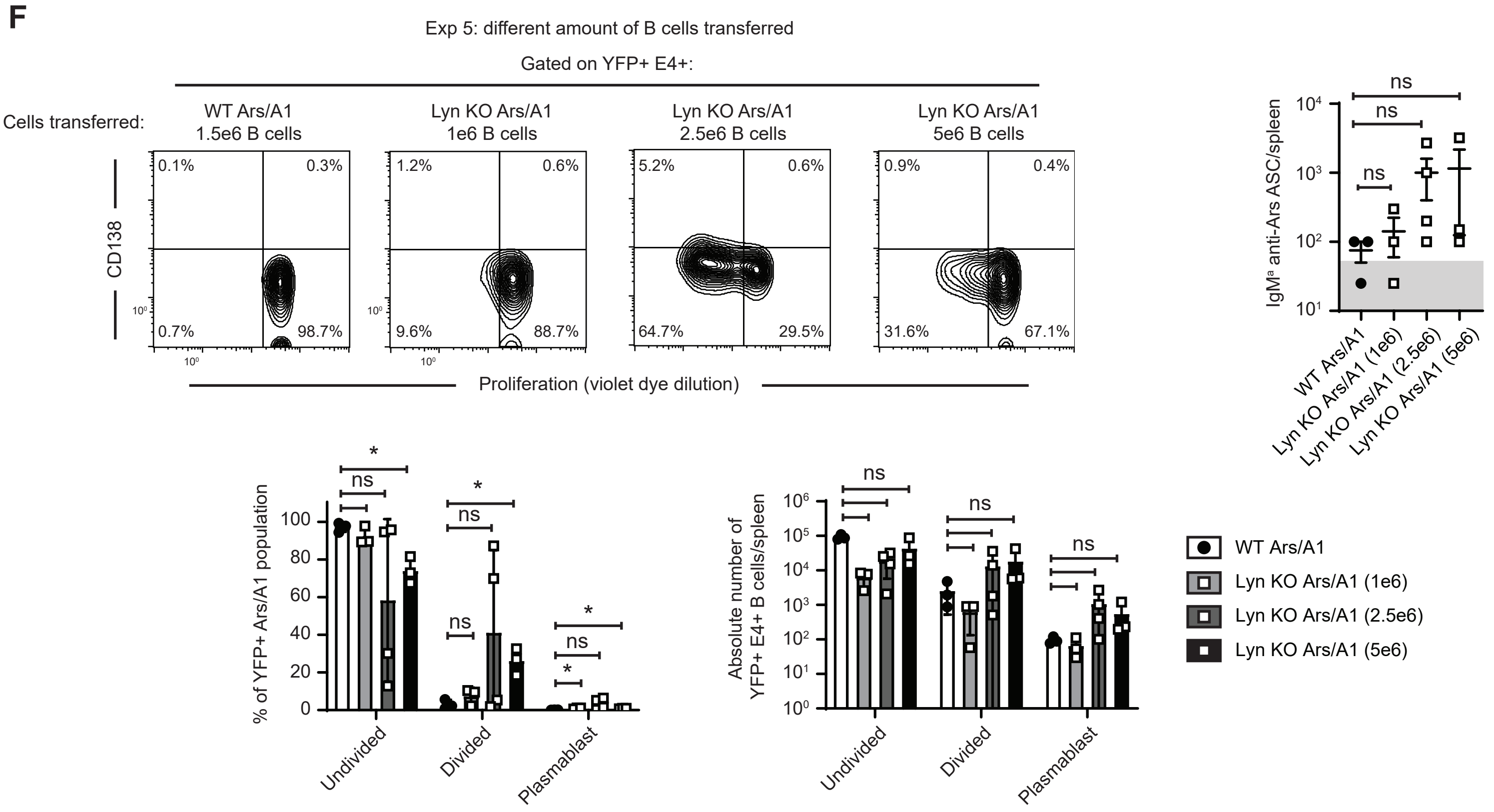


**Supplemental Figure 5. Additional data and repeat experiments for Fig 5A-F.** A-B) Additional data from the experiment shown in Fig 5A-F. A) Representative dot plots of spleens 7 or 14 days post tamoxifien treatment showing the frequency of YFP+ E4+ events. B) Absolute number of YFP+E4+ events/spleen (n=3-7 mice/genotype/timepoint). C-F) Four repeat experiments. For each experiment from left to right representative flow plots are shown to illustrate proliferation and differentiation into plasma blasts, distribution of YFP+ E4+ Ars/A1 B cells between undivided (high proliferation dye), proliferated (low proliferation dye) and plasmablasts (low proliferation dye and CD138+), absolute number of YFP+ E4+ Ars/A1 B cells among the different states, and the number of Ars/A1 IgMa antibody-secreting B cells  measured by ELISPOT. C) Experiment 2: 3e6 B cells were transferred (n=4-6).D) Experiment 3: 1.5e6 B cells were transferred (n=4-5). E) Experiment 4: 3e6 or 8e6 B cells were transferred (n=4) note that the differnt doses of Lyn B cells were from 2 different pools of donors. F) Experiment 5: 1e6, 2.5e6 or 5e6 B cells were transferred (n=3-4). Error bars represent mean ± SD. Two-tailed unpaired Student’s t test was used in the frequency graphs and a Mann-Whitney U test was used in graphs depicting absolute numbers.   ns, P > 0.05; *, P < 0.05
